# Astrocyte-secreted glypican-4 drives APOE4-dependent tau pathology

**DOI:** 10.1101/2021.07.07.451493

**Authors:** Sivaprakasam R. Saroja, Kirill Gorbachev, Julia TCW, Alison M. Goate, Ana C. Pereira

## Abstract

The aggregation of tau proteins into insoluble filaments and the spread of these filaments across brain regions are the major drivers of neurodegeneration in tauopathies, including in Alzheimer’s disease (AD). Apolipoprotein E4 (*APOE4*), a crucial genetic risk factor for late-onset AD, has been shown to exacerbate tau pathology in mouse models. However, the exact mechanisms through which APOE4 induces tau pathology remains unknown. Here, we report that the astrocyte-secreted protein glypican-4 (GPC-4), which we identify as a novel binding partner of APOE4, drives tau pathology. We discovered that GPC-4 preferentially interacts with APOE4 in comparison to APOE2, considered to be a protective allele to AD, and post-mortem APOE4-carrying AD brains highly express GPC-4 in neurotoxic astrocytes. The astrocyte-secreted GPC-4 induced both tau accumulation and propagation *in vitro*. CRISPR/dCas9 mediated activation of GPC-4 in a tauopathy mouse model robustly induced tau pathology. Using *in vitro* Tau FRET-biosensor cells, human iPSCs-derived astrocytes and an *in vivo* mouse model, we found that APOE4-induced tau pathology was greatly diminished in the absence of GPC-4. We further show that APOE4-mediated surface trafficking of APOE receptor low-density lipoprotein receptor-related protein 1 (LRP1) through GPC-4 can be a gateway to tau spreading. Together, our data comprehensively demonstrate that APOE4-induced tau pathologies are directly mediated by GPC-4.

## INTRODUCTION

AD is the most common type of dementia, accompanied by neurodegeneration and cognitive decline. The total cost of medical care for the treatment of AD in 2020 was estimated at $305 billion in the United States, and this cost is expected to increase to more than $1 trillion as the population ages^1^. There is a critical need for therapeutic drugs that may prevent the disease or slow-down the rate of disease progression. Inherited autosomal-dominant forms of AD (familial AD) are caused by the presence of mutations in amyloid precursor protein gene (*APP*) or presenilin genes (*PSEN1* and *PSEN2)*^2^. Familial AD represents less than 1% of all AD cases. The remaining AD cases do not carry such autosomal mutations, and they are termed as late-onset or sporadic AD^3^. Although the exact causes of sporadic AD remain unclear, the apolipoprotein variant E4 (*APOE4*) is known as a highest genetic risk factor for sporadic AD.

Apolipoprotein (APOE) plays a major role in the circulation of high-density and very-low-density lipoproteins and mediates the transport of lipids between the cells ^4^. Human APOE is expressed in three genetic variants: APOE2, APOE3, and APOE4. These variants differ in the position of two amino acid residues: APOE3 has a cysteine at position 112 and arginine at position 158, APOE2 has a cysteine at both positions, and APOE4 has arginine at both positions^5^. Among these three APOE isoforms, APOE4 is the most influential genetic risk factor for late-onset Alzheimer’s disease (AD). The increase in AD risk varies depending on ancestral background, sex, and multiple genetic or environmental factors. However, as a rough estimate, having a single *APOE4* allele increases AD risk 2- to 4-fold, and having two *APOE4* alleles increases AD risk approximately 8- to 16-fold ^6^. *APOE4* carriers also develop AD pathologies earlier compared to non-carriers ^6,7^. In contrast, *APOE2* carriers have a lower likelihood of developing AD; therefore, *APOE2* protects against AD^7–12^. Aβ plaques and neurofibrillary tau tangles are characteristic features of AD pathology^13,14^. AD patients with cognitive impairment often show a strong correlation with tau accumulation and spreading^15–21^. AD patients homozygous for *APOE4* suffer from significant cerebral atrophy^22–24^, and animal studies have demonstrated that pathological tau drives cerebral atrophy^25,26^. However, the mechanisms that drive tau pathology in *APOE4*_carrying individuals/animals are not well understood.

In the brain, APOE is secreted by glial cells, primarily astrocytes^6^. Cholesterol and phospholipids produced by astrocytes in the form of APOE-containing high-density lipoprotein-like particles are vital for neuronal survival^27^. Astrocytes play a critical functional role in the central nervous system, and the loss of physiological astrocytic functions can be a main contributor to neurodegeneration^28^. Recent studies suggest that a gain of toxic functions leads to astrocyte-mediated neurodegeneration in AD^29,30,31,32^. Although emerging studies suggest that neurotoxic astrocytes may be major drivers of neurodegeneration in AD cases, the molecular interconnection between astrocytes and APOE4-mediated tau pathology has yet to be resolved. Here, we report that astrocyte-secreted glypican 4(GPC-4) strongly interacts with APOE4 and exacerbates APOE4-induced tau pathology.

## RESULTS

### APOE variants and tau pathology

Tau pathology in the prefrontal cortical (PFC) regions represent the clinical dementia rate and their heterogeneity in AD patients^21,33^. Therefore, we first examined tau pathology in the PFC regions of human APOE variants. The sample details are given in the supplementary table 1. We observed the presence of neurofibrillary tangles, neuropil threads, and neuritic plaque in AD patients **(Fig. 1A)**. The neurofibrillary tangles, neuropil threads, and neuritic plaque are different forms of tau-associated neurofibrillary changes observed in the AD brain ^34^. The number of AT8 (Ser202 and Thr205) and aceTau (Lys174) positive neurons were significantly higher for *APOE4/4* compared to *APOE3/3* and *APOE2/2* AD patients **(Fig. 1B, C)**. All the subjects had amyloid-beta depositions (**Fig. S1A**). We next investigated whether APOE2 and APOE4 differentially influenced tau spreading/uptake. To address this question, we cocultured neurons from *MAPT* knockout (KO) mice that expressed GFP and from *PS19* mice (carrying the human P301S mutation in the MAPT gene) **(Fig. S1B, C)**. The tau^+^ signal in the GFP^+^-cocultured neurons from these two groups (*PS19*+*MAPT*^*KO*^) of mice is thought to be the tau protein released from *PS19* neurons. Figure 1D shows the neuronal cultures from *PS19* (left), *MAPT*^*KO*^ (middle) and both (right). Note the presence of the tau+ signal in the GFP^+^ neurons (MAPT KO neurons), suggesting that the tau proteins were released by the *PS19* neurons and were taken up by the GFP+ neurons **(Fig. 1D [right])**. To avoid any cross-reactivity during imaging, monocultures of *PS19* and *MAPT*^*KO*^ were used as baseline signals of GFP and AT8, respectively, for the cocultures (*PS19*+*MAPT*^*KO*^). We isolated APOE2 and APOE4 particles from frozen PFC regions of normal human *APOE2/2* and *APOE4/4* carrier brains **(Fig. S1D)**. When the neuronal cultures were treated with APOE isoforms, we observed that APOE4 robustly enhanced tau spreading/uptake **(Fig. 1E, F)**. Upon adding APOE2 with APOE4 to the neuronal cultures, we observed no APOE4-mediated increase in tau spreading/uptake **(Fig. 1E, F)**. Further, we found that APOE4 induced significantly more AT8 levels in *PS19* neuronal culture, but APOE4-induced AT8 levels were reduced in the presence of APOE2 **(Fig.S1E-G)**.

**Figure 1.**
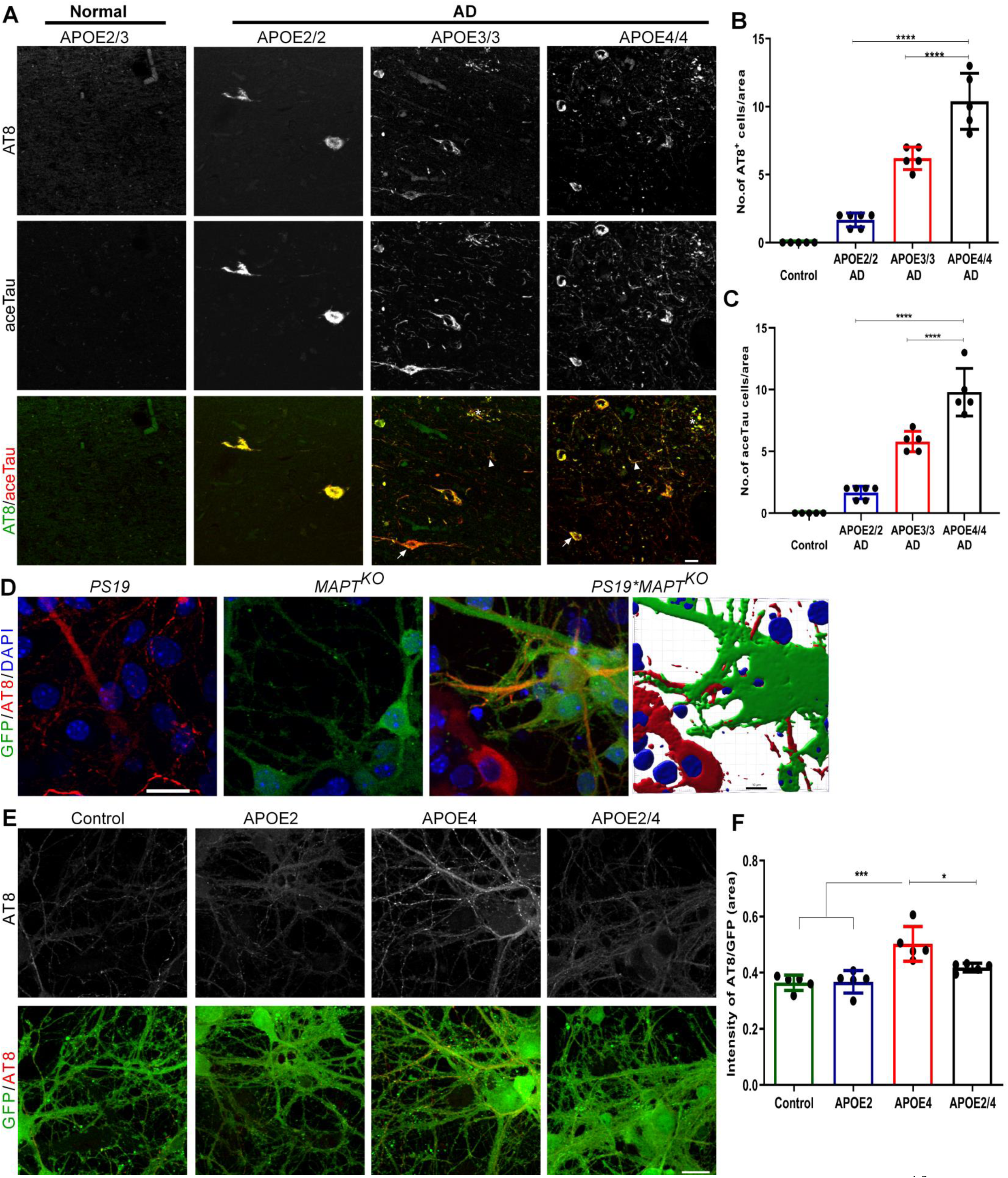
The brains of postmortem APOE4 AD patients accumulate more tau proteins and APOE4 enhances tau propagation/uptake. **A**) Representative IHC images of postmortem PFC tissues from *APOE2/3*(control), *APOE2/2*(AD), *APOE3/3* (AD) and *APOE4/4* (AD) individuals stained with AT8 and aceTau (Lys174) antibodies (n=5-6) show the presence of Neurofibrillary tangles (arrow), neuropil threads (arrowhead) and neuritic plaque (asterisk). **B, C**) Compared to *APOE3/3* and *APOE2/2, APOE4/4* carriers show the presence of more neurofibrillary tangle-containing neurons. **D**) Neuronal monocultures from *PS19* (left), *MAPT*^*KO*^ (middle), and mixed neuronal culture of *PS19*\**MAPT*^*KO*^ (right) showing the expression of tau (AT8), GFP and uptake of tau proteins, respectively. *PS19*\**MAPT*^*KO*^ neurons were 3D-reconstructed using imaris software. **E**) We treated the mixed neurons (*PS19*\**MAPT*^*KO*^) with APOE2 or APOE4 particles, and accessed tau pathology after 4 days. Representative IHC staining of APOE treated neuronal culture shows that APOE 4 treatments enhanced the tau spreading whereas APOE2 significantly reduced APOE4-mediated tau spreading (**F**). n=5, one-way ANOVA, *P<0.05, ***P<0.001 and ****P<0.0001. IHC scale bars=20 µm.

### Interlink between GPC-4 and APOE4

Alboleda-Velasquez *et al* recently reported that an autosomal mutation carrying-AD patient did not develop severe cognitive impairments until her seventies. She also had an unusual mutation in the *APOE3* gene (Christchurch R136S mutation)^16^. Compared to purified human APOE3 proteins, APOE3 protein with R136S mutation had a weaker interaction with heparin^16^. It is proposed that these distinct interactions of APOE R136S with heparin may be a reason that the previously mentioned patient did not develop dementia and had less tau spreading compared to her counterparts^16^. We therefore reasoned that Heparan Sulfate Proteoglycans (HSPGs) may be involved in APOE4-mediated tau pathology. We first screened a list of HSPGs and their interactions with APOE2 or APOE4, and found that glypican-4 (GPC-4) strongly binds with APOE4 compared to APOE2 (**Fig. 2A-C and S2A**). We incubated purified human GPC-4 protein either with purified human APOE2 or APOE4 protein at room temperature for 1 h, as described in Figure 2A, and analyzed the samples with a native gel. The APOE2+GPC-4 combination did not show any major shifts, whereas the APOE4+GPC-4 mix showed a robust shift with both GPC-4 and APOE antibodies. Furthermore, treatment with 2-mercaptoethanol disturbed the shift of GPC-4+APOE4, suggesting that GPC-4 and APOE4 are in direct interaction (**Fig. 2A**). We next validated this finding in postmortem human brain tissues. We co-immunoprecipitated GPC-4 proteins from *APOE2/2* and *APOE4/4* human brains using APOE antibodies (**Fig. 2B**). The levels of eluted GPC-4 proteins were normalized by corresponding eluted APOE proteins. **Figure 2C** shows that GPC-4 preferentially binds with APOE4 *in vivo*.

**Figure 2.**
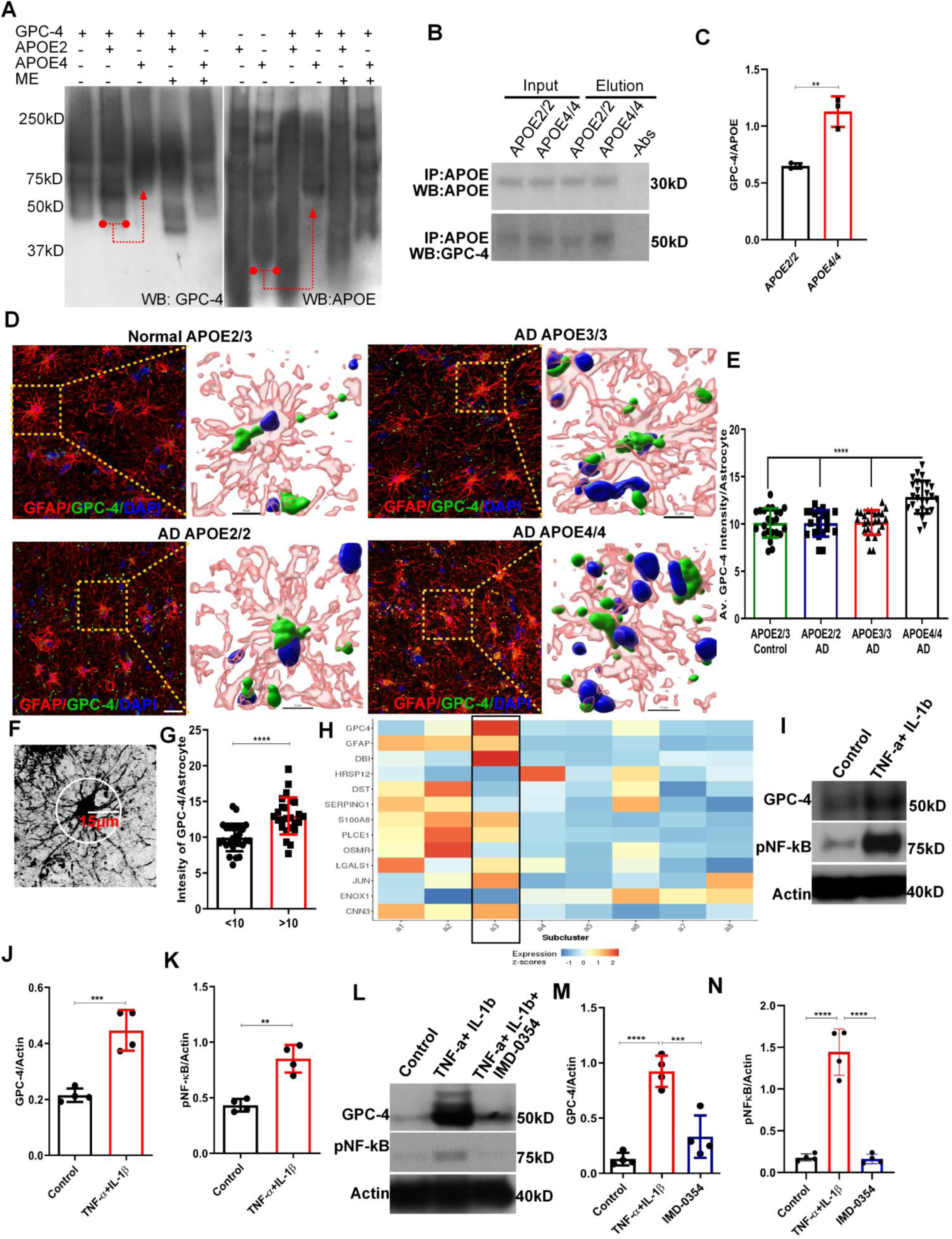
APOE4 interacts with GPC-4 and APOE4 AD patients express more GPC-4 in neurotoxic astrocytes. **A)** GPC-4 protein was incubated with either APOE2 or APOE4 proteins at room temperature for 1h, and then separated them by a native gel. Western blot analysis with GPC-4 (left panel) and APOE (right panels) antibodies revealed that combinations of APOE4+GPC-4 were shifted while no observable changes occurred with APOE2+GPC-4. A red arrow in the left panel indicates GPC-4 proteins were shifted when combined with APOE4, but not with APOE2. A red arrow in the right panel indicates APOE4 proteins were shifted when combined with GPC-4, whereas APOE2 proteins were not shifted when combined with GPC-4. ME=β-mercaptoethanol. **B, C)** Proteins isolated from *APOE2/2* and *APOE4/4* human brains were co-immunoprecipitated with APOE antibodies (n=3) (B). The levels of immunoprecipitated GPC-4 proteins were normalized with corresponding APOE immunoreactive bands (C). It shows that the APOE/GPC-4 complex is significantly higher in *APOE4/4* compared to *APOE2/2*. **D, E)** Representative IHC staining of *APOE2/3*(control), *APOE2/2*(AD), *APOE3/3* (AD) and *APOE4/4* (AD) tissues with GFAP and GPC-4 antibodies show that *APOE4* carrying AD patients expressed significantly higher levels of GPC-4 in astrocytes compared to control and other APOE genotypes (E). n=5-6. **F, G)** Astrocytes from *APOE4/4* AD brains were grouped into two categories based on the number of the branches at 15μm radius (F). Group 1: less than 10 branches. Group 2: more than 10 branches. Astrocytes with more branches expressed significantly elevated levels of GPC-4 (G). **H)** AD-associated astrocytic genes were selected from Habib *et al, Nature Neuroscience*, 2020 and generated a heat map with subtypes of astrocytes from Grubman *et al, Nature Neuroscience*, 2019. Disease-associated genes were enriched in subtypes 2 and 3, and GPC-4 was enriched in a subtype 3. **I-K)** Western blot analysis from astrocyte culture shows that treatment with TNF-α and IL-1β significantly increased expression of GPC-4, and it also activated the NF-κB pathway. **L-N**) Western blot analysis from astrocyte culture treated with NF-κB pathway blocker, IMD-0354, reversed TNF-α and IL-1β induced expression of GPC-4. n=4. one-way ANOVA or unpaired t-test, **P<0.01, ***P<0.001 and ****P<0.0001. IHC scale bars=20 µm.

We next examined whether GPC-4 was differentially expressed in human APOE variants. Confirming previous research^35,36^, we observed that GPC-4 is specifically expressed in astrocytes (**Fig. S3A**)^37^. IHC staining showed that APOE4-carrying AD patients expressed more GPC-4 protein in astrocytes than non APOE4-carrying AD patients **(Fig. 2D, E)**. Astrocytes represent a diverse population of cells with varying complex morphologies and functions^38,39^. Astrocytes with fewer branches are classified as resting astrocytes, while astrocytes with more branches are classified as disease-associated astrocytes (neurotoxic)^40,32,29^. Figures 2F and 2G show that astrocytes with more branches express significantly higher levels of GPC-4 protein, suggesting that neurotoxic astrocytes express GPC-4. We analyzed previously published single-cell RNA-sequencing (ScRNAseq) studies from human and mouse brains to independently validate these results. The scRNAseq studies revealed the presence of AD-associated genes and astrocytes in mice and humans ^37,41,42^. We generated a heatmap of AD-associated genes to test which subtypes of astrocytes express GPC-4 (**Fig. 2H**). AD-associated genes are mainly expressed in astrocyte subtypes 2 and 3. Interestingly, GPC-4 was expressed within AD-associated astrocyte subcluster 3 (**Fig. 2H**). It suggests that GPC-4 is expressed in AD-associated A1-reactive or neurotoxic astrocytes. A1-reactive astrocytes, which are induced by microglial factors such as TNF-α and IL-1β, are considered as neurotoxic astrocytes^43^. TNF-α and IL-1β treatment in astrocytes induced significantly higher levels of GPC-4 protein **(Fig. 2I-K)**. Furthermore, TNF-α and IL-1β-induced GPC-4 expression was blocked in the presence of NF-κB inhibitor IMD-0354 **(Fig. 2L-N)**, suggesting that the NF-κB dependent pathway regulates expression of GPC-4 in disease-associated or neurotoxic astrocytes.

### Glypican-4 induces tau pathology *in vitro* and *in vivo*

We next asked whether GPC-4 plays a role in tau pathology. We treated the *PS19*-derived neurons with purified human GPC-4 and found that it robustly induced AT8 and PHF1 (paired helical filament) levels (**Fig. 3A-D**). The IHC experiments confirmed these findings (**Fig. 3E, F**). We next treated a neuronal culture with astrocytes-conditioned medium (ACM), as described in Figures 3G (Fig. S3B). Like purified human GPC-4 protein, ACM-treated neurons expressed significantly higher levels of AT8 (**Fig. 3H-J**). We next treated the astrocytes with GPC-4 shRNA **(Fig.S3C, D)**, and collected GPC-4 deprived ACM (**Fig. 3K**). Interestingly, GPC-4 deprived ACM failed to induce tau pathology **(Fig. 3L-N)**. We cocultured neurons of *PS19* and *MAPT*^*KO*^ animals to test the effect of GPC-4 on tau spreading/uptake, and found that GPC-4 increased tau spreading from *PS19* neurons to *MAPT*^*KO*^ neurons (**Fig. 3O, P**).

**Figure 3.**
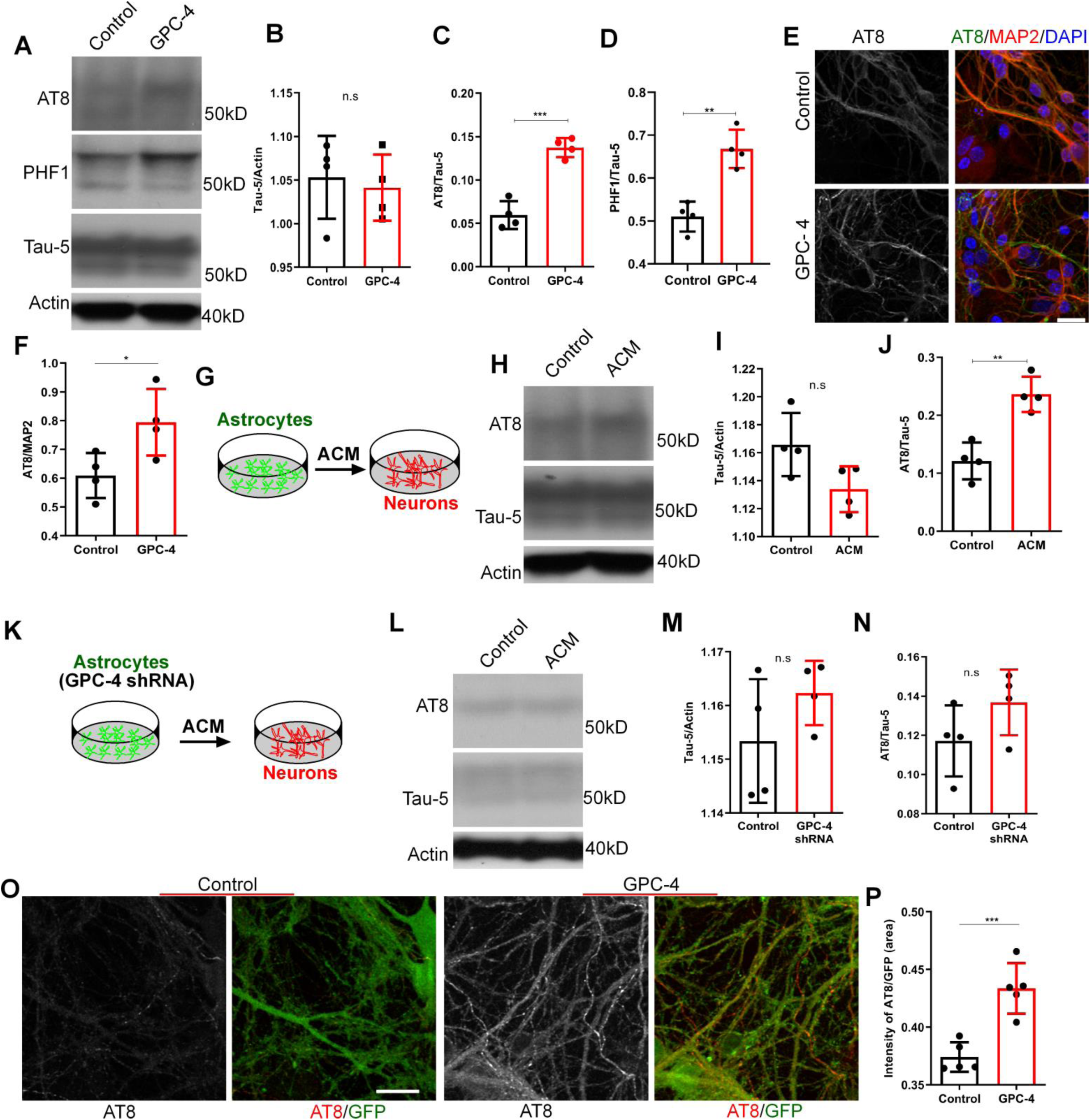
GPC-4 induces tau pathology *in vitro*. **A-D)** Western blot analysis of proteins isolated from neuronal culture, which were treated with GPC-4 protein, shows that GPC-4 significantly enhanced pTau levels (C, D) whereas no changes were observed in total tau protein (B). **E, F)** Representative IHC staining of GPC-4 treated neuronal culture with AT8 and MAP2 antibodies demonstrates that GPC-4 treatment enhanced pTau levels (E). **G)** Schematic diagram shows that WT astrocyte-conditioned media (ACM) from astrocyte culture was added to *PS19* neuronal culture. **H-J)** Addition of ACM to neuronal culture increased the pTau (AT8) levels whereas total tau proteins were unaltered. **K)** Schematic diagram shows that astrocytes were treated with GPC-4 shRNA and the resulting GPC-4 deprived ACM was added to neuronal culture. **L-N)** Addition of GPC-4 deprived ACM failed to induce tau phosphorylation in neurons. **O, P)** Representative IHC staining of GPC-4 treated neuronal co-culture (*PS19* MAPT*^*KO*^ animals) shows that GPC-4 treatment enhanced tau spreading/uptake. n=4-5, unpaired t-test, IHC scale bars=20 µm. *P<0.05, **P<0.01, ***P<0.001 and ****P<0.0001.

We next sought to investigate a possible link between GPC-4 and tau pathology *in vivo*. An adeno-associated virus that encodes eGFP-2PA-hTauP301L mRNA was recently used to study tau propagation from one neuron to another ^44,45^. 2PA sites will cause the translation of independent eGFP and hTauP301L proteins. While the donor (initially infected) neurons will express both eGFP and hTauP301L proteins, the released hTauP301L proteins will be taken up by synaptically connected neurons (the recipients). We injected eGFP-2PA-hTauP301L AAV virus in the ipsilateral CA1 regions and control/GPC-4 shRNA in the contralateral CA1 regions **(Fig. S4A)**, and examined the samples after 4 weeks to investigate the effects of GPC-4 in tau spreading. As expected, we observed a robust expression of eGFP and hTauP301L in the ipsilateral sites (**Fig. 4A**). We also observed the propagation of hTauP301L proteins to the contralateral CA1 regions (**Fig. 4A**). Interestingly, GPC-4 shRNA reduced the levels of hTauP301L in contralateral CA1 regions (**Fig. 4B**). This finding validates our *in vitro* data **(Fig.3O, P)**. We next induced the expression of GPC-4 in *PS19* animals using a CRISPR/dCas9 system to investigate the role of GPC-4 in tau accumulation *in vivo*. After one week of incubation, we observed a tremendous expression of GPC-4 proteins in astrocytes **(Fig. 4C, D)**, but not in neurons **(Fig. S4B)**. Following 3 weeks of induction, we observed a robust phosphorylated tau protein in the hippocampal region **(Fig. 4E, F and S4C)**. Additionally, induction of GPC-4 in the cortical regions also induced tau accumulation **(Fig. 4G, H)**.

**Figure 4.**
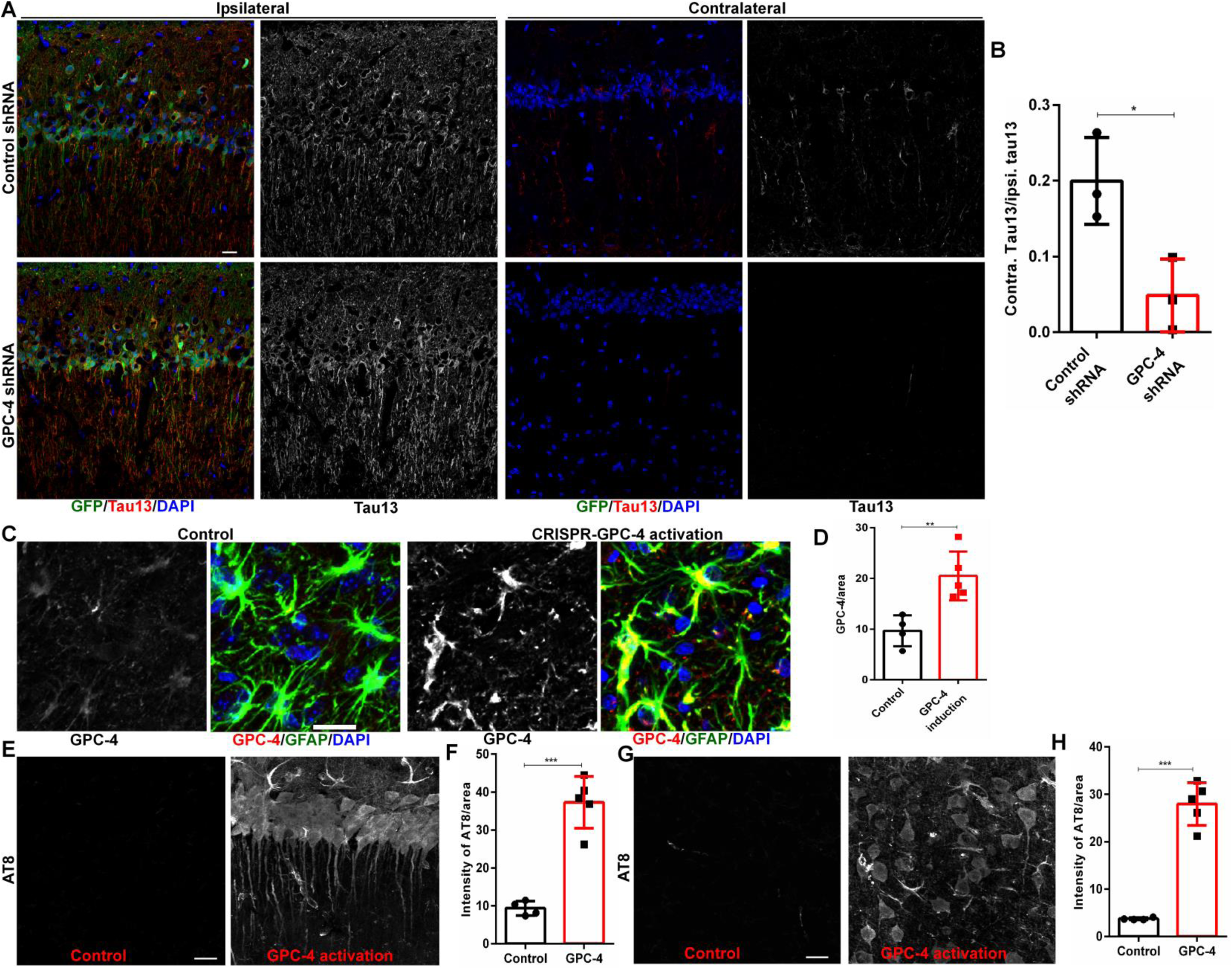
GPC-4 induces tau pathology *in vivo*. **A, B)** we injected eGFP-2PA-hTauP301L AAV virus in the ipsilateral CA1 regions and control/GPC-4 shRNA in the contralateral CA1 regions, and examined them after 4 weeks. Representative IHC images with GFP and human tau specific antibody Tau13 show that GPC-4 shRNA treatment significantly reduced tau spreading in the contralateral CA1 regions. **C, D)** In order to induce the expression of GPC-4 proteins, we activated the GPC-4 gene by injecting GPC-4 CRISPR/dCas9 lentivirus activation systems in the cortex or hippocampus. Representative IHC with GPC-4 and GFAP antibodies show that, following a one week of injection, GPC-4 CRISPR/dCas9 robustly induced GPC-4 expression compared to control lentiviral activation particles. **E, F)**. IHC images with the AT8 antibody show that GPC-4 induced significantly higher levels of pTau in CA1 regions of the hippocampus. **G, H)**. IHC images with AT8 antibody shows that GPC-4 induced significantly higher levels of pTau in the cortex. n=4-5, unpaired t-test, *P<0.05, **P<0.01 and ***P<0.001. IHC scale bars=20 µm.

### Glypican-4 is necessary for APOE4-mediated tau propagation

Given that GPC-4 induced tau pathology and interacted with APOE4 (Figs. 2-4), we reasoned that GPC-4 would play an important role in APOE4-mediated tau pathology. We treated the primary neuronal culture (WT) with ACM alone, APOE4+ACM, and APOE4 with GPC-4 deprived ACM for 24 h and then incubated it with 1 µg/ml of human tau protein (#842501, labome) for 1 h. After washing, IHC with human tau antibody HT-7 showed that APOE4 increased tau uptake, but this increase was reversed in the absence of GPC-4 (**Fig. 5A, B**). This result indicates that GPC-4 regulates APOE4-induced tau uptake. We next examined a possible interaction between GPC-4 and APOE4 in tau aggregation *in vitro*. We used Tau FRET-biosensor cells to monitor tau protein aggregation^46^. Human AD-derived insoluble tau protein treatment caused aggregation of endogenous tau protein, confirming the prion-like nature of our isolated tau proteins **(Figs. S5A, 5C)**. Interestingly, APOE4 treatment increased tau aggregation, and the absence of GPC-4 diminished APOE4-induced tau aggregation (**Fig. 5C, D and S5B, C**). We next investigated the role of GPC-4 in APOE4-induced tau pathology *in vivo*. We injected APOE particles in the CA1 regions of the *PS19* animals and examined this group after 3 weeks. As expected, APOE4 robustly induced tau accumulation **(Fig. 5E-H)**. Interestingly, APOE4-mediated tau pathology was dramatically reduced in the absence of GPC-4 **(Fig. 5E-H)**. We performed an additional experiment to validate the role of GPC-4 in APOE4-mediated tau pathology in human induced pluripotent stem cells (iPSCs)-derived astrocytes. Human APOE4 iPSCs cells were differentiated into mature astrocytes as previously described^47,48^**(Fig. 5I)**. The astrocytes were treated with GPC-4 shRNA **(Fig. S5D, E)**, and ACM was collected and added to *PS19*-derived neurons as described in Figure 3K. GPC-4 shRNA treatment did not show significant changes in the APOE levels of iPSCs-derived astrocytes **(Fig. S5F, G)**. While the ACM treatment from *APOE4* increased tau pathology, the GPC-4 shRNA treatment hampered the ACM-induced tau pathology **(Fig. 5J-M)**.

**Figure 5.**
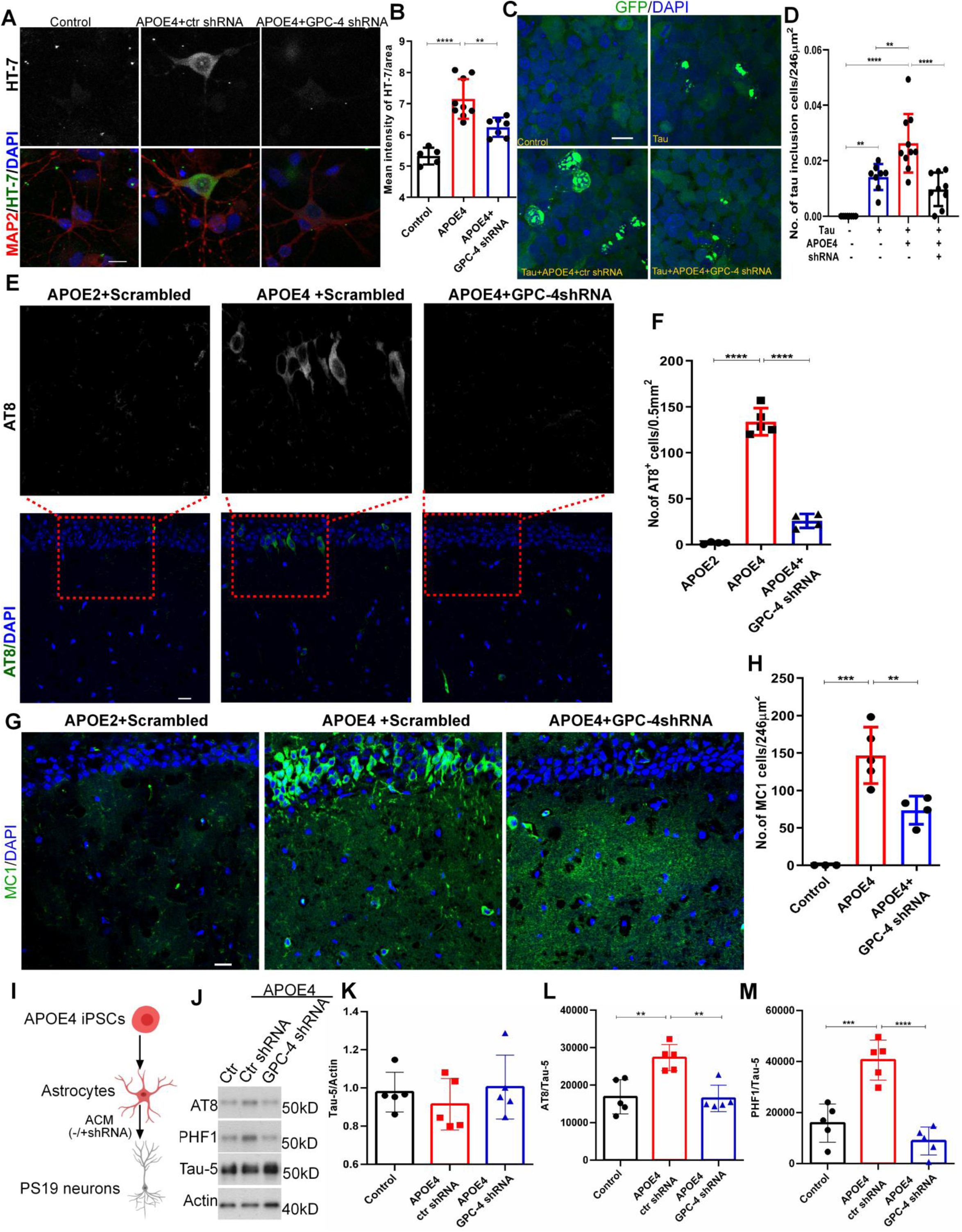
GPC-4 drives APOE4-mediated tau pathology. **A, B)** Cultured neurons (WT) were treated with ACM alone, ACM with APOE4 and GPC-4 deprived (shRNA treated) ACM with APOE4. After 24hrs, neurons were incubated with purified human tau proteins for 1hr, washed several times, and immunostained with human tau antibody (HT-7) to detect human tau proteins. Representative IHC images of APOE4 + ACM treated neuronal culture shows that APOE4 treatment enhanced tau uptake, but in the absence of GPC-4 (GPC-4 shRNA treated ACM) APOE4-induced tau uptake was significantly reduced (B). **C, D)** we used Tau FRET-biosensor cells to monitor seeding activity of AD-tau proteins in the presence of APOE4 and GPC-4. We incubated Tau FRET-biosensor cells with 0.6μg/ml tau proteins and 1μg/ml APOE4 with/without GPC-4 shRNA for 2 days, and then counted the number of tau inclusion containing cells/area. Compared to tau alone, tau with APOE4 induced significantly higher numbers of tau inclusions in Tau FRET Biosensor cells. Interestingly, APOE4-mediated tau inclusions were reduced in the presence of GPC-4 shRNA. n=8-10. **E, F)** we injected APOE2 or APOE4 particles isolated from corresponding human brains, in the absence or presence of GPC-4 shRNA. Following 3 weeks of injections, no tau pathology (AT8) was detected with APOE2. APOE4 robustly induced tau pathology, but APOE4 failed to induce tau pathology in the absence of GPC-4 (D). **G, H**) IHC staining with MC1 antibodies show that APOE4-mediated tau pathology was diminished in the presence of GPC-4 shRNA. **I-M)** Human *APOE4* iPSCs were differentiated into astrocytes, and the ACM was collected after 3 days of shRNA treatments. The *PS19* neurons were treated with ACM for 4 days. Western blot analysis shows that ACM significantly increased AT8 and PHF1 levels, and the ACM-induced tau pathologies were reduced in the absence of GPC-4. n=4-5, one-way ANOVA, **P<0.01, ***P<0.001 and ****P<0.0001. IHC scale bars=20 µm.

### Glypican -4 regulates APOE4-mediated membrane trafficking of the LRP1 receptor

Finally, we wanted to understand the molecular interaction between APOE4 and GPC4 in inducing tau pathology. The LRP1 (low-density lipoprotein receptor-related protein 1) is a major APOE receptor, and it has been suggested that it is involved in tau uptake and spreading ^44^. We therefore examined whether the APOE2 and APOE4 have differential effects on the LRP1 receptor. We monitored the effects of APOE2 and APOE4 on the total and surface LRP1 levels in WT neuronal cultures. We found that the addition of APOE2 had no effect on either the total or surface LRP1 levels (**Fig. 6A-C**). In contrast, APOE4 enhanced the trafficking of surface LRP1 levels (**Fig. 6A-C**). Interestingly, APOE4-mediated surface trafficking of LRP1 was blocked by APOE2 (**Fig. 6A-C**). Furthermore, the reduction of APOE4-induced surface LRP1 by an exocytic pathway blocker, Exo-1, suggests that the active exocytic pathways are required for APOE4-induced surface trafficking of LRP1 (**Fig. 6D-F**).

**Figure 6.**
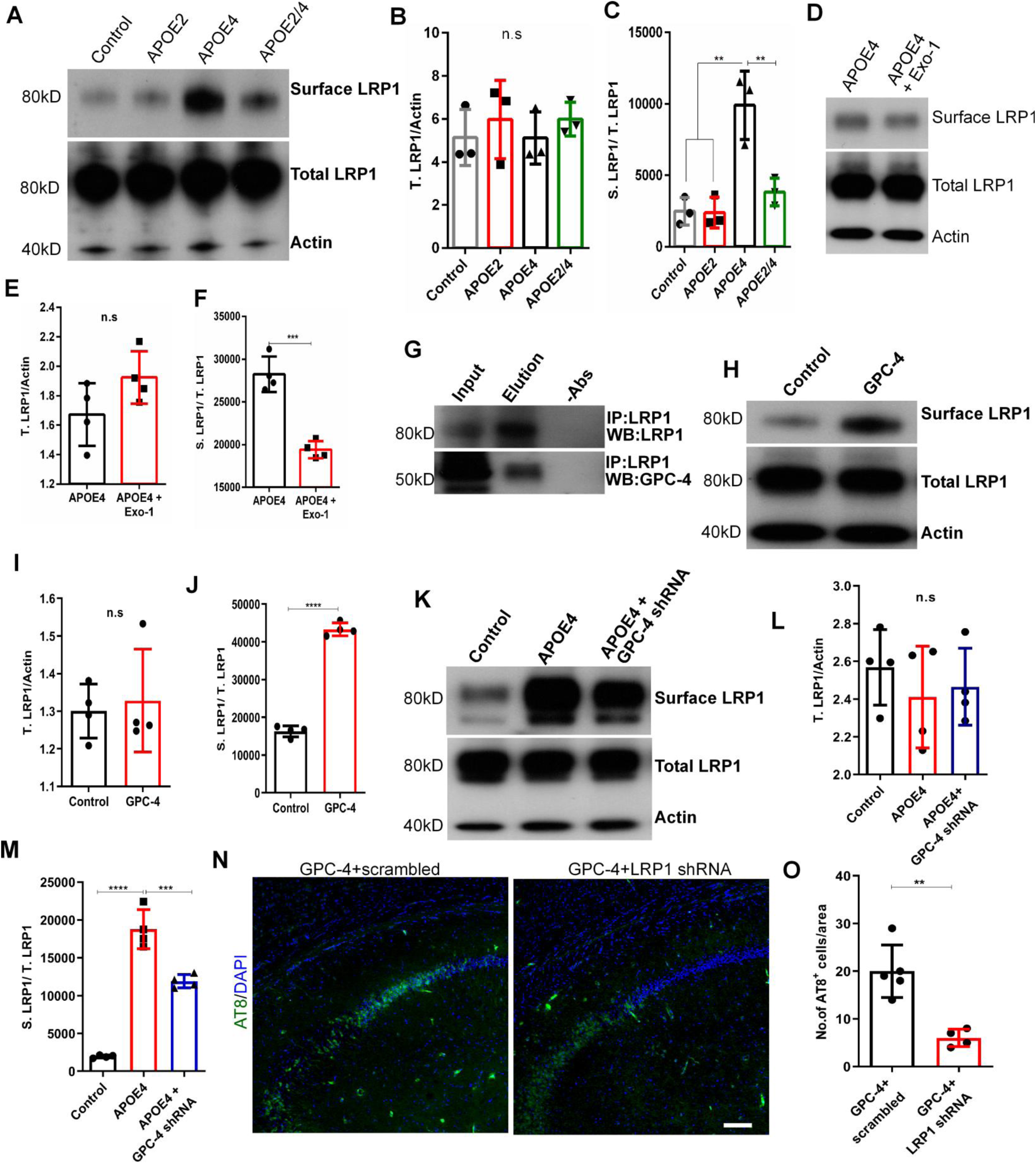
GPC-4 regulates trafficking of APOE receptor LRP1. **A-C)** Western blot analysis from neuronal culture treated with APOE (s) shows that APOE4 significantly enhanced the surface LRP1 (S.LRP1) (C). There were no changes in total LRP1 (T.LRP1) (B). **D-F)** Western blot analysis from neuronal culture treated with APOE4 or APOE4 with exocytosis inhibitor, Exo-1, show that active exocytosis is required for APOE4-mediated surface trafficking of LRP1. **G)** Co-immunoprecipitation of human postmortem brain protein samples with LRP1 antibodies and subsequent western blotting revealed that LRP1 and GPC-4 are in the same complex. **H-J)** Western blot analysis from neuronal culture treated with GPC-4 protein shows that GPC-4 significantly enhanced trafficking of surface LRP1 levels (J), whereas total LRP1 levels were not affected (I). **K-M**) Western blot analysis from neuronal culture treated with ACM alone (control), APOE4 + ACM, and with APOE4 + GPC-4 shRNA treated ACM show that APOE4 induced surface expression of LRP1 is mediated through GPC-4. The GPC-4 shRNA treatment reduced the surface expression of LRP1 in the presence of APOE4 (J). **N, O)** Representative IHC images with AT8 antibodies show that GPC-4 induced tau pathology was reduced in the absence of LRP1. n=4-5, one-way ANOVA or unpaired t-test, **P<0.01, ***P<0.001 and ****P<0.0001. IHC scale bars=20 µm.

We next reasoned that since GPC-4 is an astrocyte-secretory factor and an APOE4 binding partner, it would interact with neuronal LRP1. We first examined the likelihood of the GPC-4–LRP1 interaction to test this notion. A co-immunoprecipitation assay from human postmortem brain tissue revealed that GPC-4 directly interacts with LRP1 **(Fig. 6G)**. In the WT neuronal culture, the addition of GPC-4 had no effect on the total LRP1, whereas the surface LRP1 levels increased greatly **(Fig. 6H-J)**. We next investigated whether APOE4 is dependent on GPC-4 to induce surface LRP1 levels. We treated neurons with either APOE4 with ACM (WT mouse astrocytic culture) or APOE4 with GPC-4 shRNA treated ACM, as shown in Figure 3G and 2K. APOE4 in the absence of GPC-4 showed a significant reduction in surface trafficking of LRP1 compared to APOE4 with ACM **(Fig. 6K-M)**. Additionally, in the absence of LRP1 (LRP1 shRNA), GPC-4 induced tau pathology was significantly reduced in the CA1 regions of *PS19* mouse **(Fig. 6N, O and S6)**.

## DISCUSSION

In addition to an earlier observation that suggested the presence of a strong correlation between tau pathology and dementia^21^, the recent studies have overwhelmingly demonstrated that tau accumulation/spreading is associated with neurodegeneration and dementia^16,20,25^. Tau-based therapy has recently become attractive for clinical trials. At this juncture, it is critical to understand every step-in tau pathology. Human *APOE4* transgene in a *PS19* mouse model displayed more tau accumulation ^25,49^, and human tau PET studies in *APOE*4 carriers agree with these observations ^12,50,51^. Nevertheless, the mechanism through which APOE4 induces tau pathology remains unknown. Here, we found that the astrocytic protein GPC-4 preferably interacts with APOE4, and the brains of postmortem APOE4 AD patients highly expressed GPC-4 in neurotoxic astrocytes. We showed that GPC-4 induced tau accumulation and propagation *in vitro*. CRISPR/dCAS9-mediated activation of GPC-4 induced tau pathology *in vivo*. In the absence of GPC-4, APOE4-mediated tau pathology was greatly diminished in Tau FRET biosensor cells and *PS19* neurons. We further demonstrated a molecular interlink between GPC-4, APOE4, and its receptor LRP1, which is also implicated in tau pathology.

GPC-4 is one of the six members of the glypican family. GPC-4 is an astrocyte-secreted protein that has been shown to regulate synaptic plasticity in the developing brain^36,35^. We found that GPC-4 preferentially binds with the APOE4 isoform, a risk allele to developing AD over the APOE2 isoform, a protective allele to AD. A subtype of activated A1 astrocytes are considered neurotoxic astrocytes, whose secretory molecules may be involved in the worsening of AD pathology^43,52^. Our data showed that GPC-4 is mainly expressed within a neurotoxic astrocytic population in APOE4 carrying AD patients. This result is consistent with the scRNAseq analysis of AD patients and AD mouse models^37,53^. We also found that the expression of GPC-4 was regulated via the NF-kB pathway in the presence of microglial factors. In fact, a mouse study showed that human APOE4-expressing mice bred with *PS19* tau mouse model displayed increased microglial activity^25^.

The role of HSPGs in Aβ pathology is well documented^54^. It has been shown that HSPGs are involved in the early stage of Aβ pathology in AD^55–57^. However, the role of HSPGs in tau pathology is poorly understood. An AD patient with an autosomal dominant mutation who had not developed cognitive impairments for several decades before expected had a mutation in the *APOE3* allele (R136S, Christchurch)^16^. While normal APOE3 strongly interacted with HSPGs *in vitro, APOE3* R136S showed a weak interaction^16^. This suggests that HSPGs can potentially have an important impact on tau pathology. After demonstrating that GPC-4 preferentially binds with APOE4 over APOE2, we showed that GPC-4 enhances the phosphorylation of tau proteins and their propagation *in vitro*. Further activation of GPC-4 *in vivo* using a CRISPR/dCas9 system induced tau pathology in PS19 animals. Given that mice do not express APOE isoforms, injection of APOE particles isolated from the human brain more closely resembles human conditions, enabling the study of the effect of their role in AD pathologies. Interestingly, APOE4-induced tau pathology was greatly diminished in the absence of GPC-4, suggesting that GPC-4 plays a critical role in tau pathology. It has been shown that GPC-4 enhances neuronal excitability^36^. Similarly, it is also known that APOE4-carrying AD patients/animals display hyperexcitability^58,59^. Given that neuronal activity is proposed to enhance tau propagation^60^, future studies are warranted to examine the interlink between neuronal activity and GPC-4/APOE4 in tau pathology.

It has been recently proposed that the APOE receptor LRP1 binds with tau proteins and internalizes them^44^. We showed that APOE4 induces cellular surface trafficking of LRP1 and tau pathology. Notably, the addition of APOE2 did not alter the surface levels of LRP1 and tau pathology, instead, it attenuated the APOE4-induced trafficking of LRP1 and tau pathology. These results suggest that APOE2 is involved in rescue mechanisms in the presence of APOE4. We further showed that GPC-4 is in the complex with LRP1 and APOE4, and that APOE4-mediated surface trafficking of LRP1 is likely dependent on GPC-4. This differential action of APOE isoforms on the surface trafficking of LRP1 thereby likely relies on the presence of associated factors, such as GPC-4, which interacts more strongly with APOE4 than APOE2.

A limitation of our study, in fact in the AD field itself, is the lack of animal models for sporadic AD. As a result, researchers utilize either dominant mutations containing Aβ animal models (such as *5xFAD*) or frontotemporal dementia tau animal models (such as *PS19*) to understand Aβ and tau pathology, respectively^25,61,62^. However, in sporadic AD, tau and Aβ accumulate without mutations in the MAPT or APP/PSEN genes. The lack of genetically modifiable animal models to study sporadic AD has been a major setback for the field. Therefore, while other models (including injection of AD brain -derived tau species) have dramatically improved our understanding of tau pathology^63^, modeling human sporadic AD-related organisms or organoids will likely improve therapeutic options for AD patients. Further, our *in vitro* studies showed that APOE2 reverses APOE4-induced tau pathology and surface trafficking of LRP1 receptors. However, our studies did not address how APOE2 reverses APOE4-mediated effects. Furthermore, most of current studies compare individual APOE variants. The heterozygous APOE carriers represent the majority of AD cases compared to homozygous APOE carriers^6,64^. Therefore, understanding how APOE4-mediated activity is affected in the presence of other APOE isoforms, such as APOE2, is important. In conclusion, we propose that GPC-4 works along with APOE4 and its receptor LRP1 to induce tau pathology, and targeting GPC4/APOE4/LRP1 complexes can open therapeutic windows against tau pathology in AD.

## Acknowledgments

This work was supported by funding from NIH R01 AG063819 (ACP), NIH R01AG064020 (ACP), Paul B. Beeson Emerging Leaders Career Development Award in Aging K76 AG054772 (ACP), the Bright Focus Foundation (ACP), the DANA Foundation (ACP), the Alzheimer’s Drug Discovery Foundation (ACP), the Alzheimer’s Association (ACP), Carolyn and Eugene Mercy Research Gift (ACP), Karen Strauss Cook Research Scholar Award (ACP), NIH K01AG062683 (JTCW), U01AG058635 (AMG) and U19AG069701 (JTCW, AMG). The content is solely the responsibility of the authors and does not necessarily represent the official views of the National Institutes of Health. We thank Mount Sinai Brain Bank and Banner Sun Health Research Institute for providing human brain samples, Patrick Hof, Diede Broekaart, Saraswathi Subramaniyan and Sam Gandy for insightful comments. We Neeva Shafiian for assisting with quantification analysis.

The authors have no conflict of interest.

## Figure legends

**Supplementary Figure 1.**
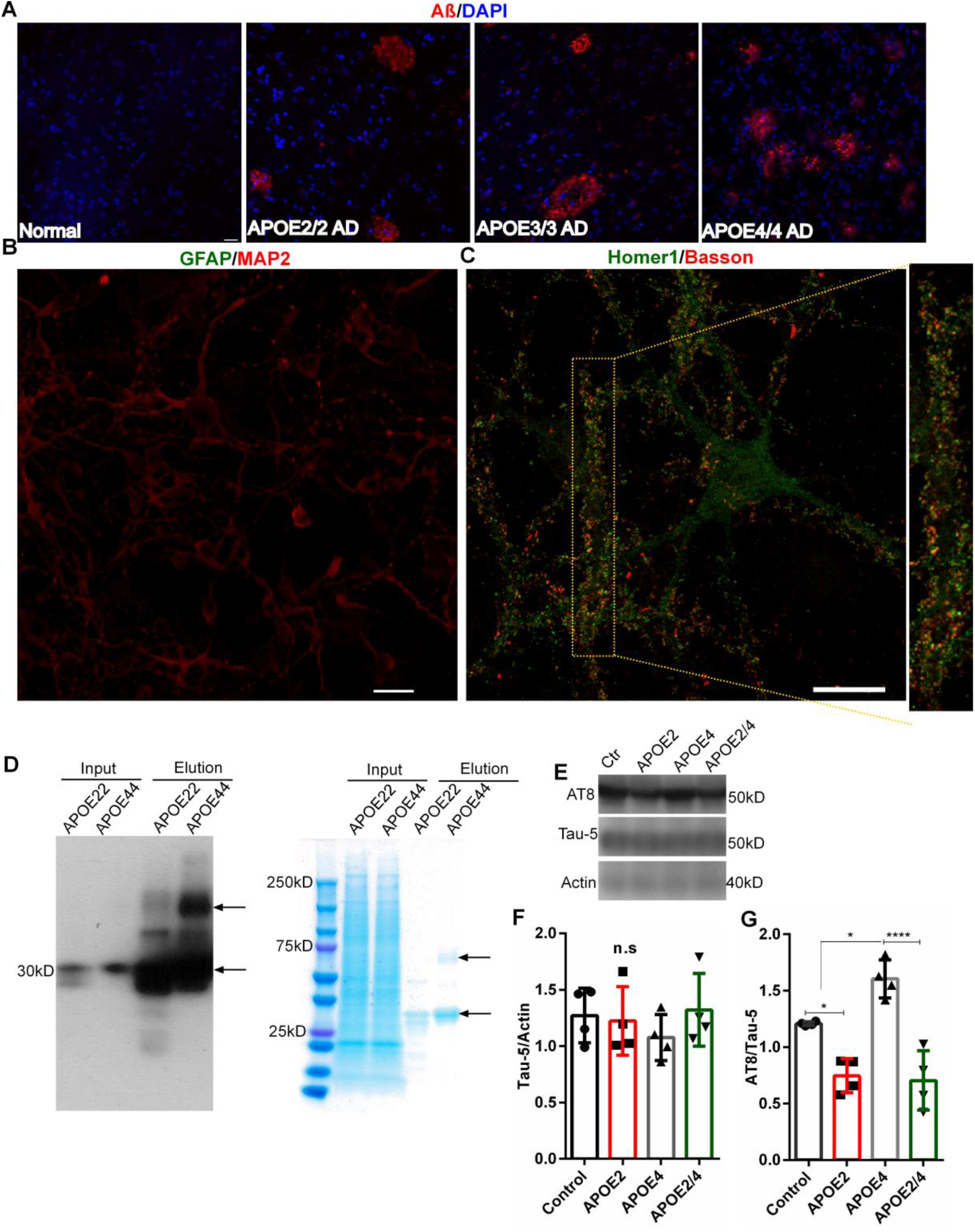
**A)** Representative IHC images from APOE2/2, APOE3/3 and APOE4/4 AD cases show the presence of beta-amyloid depositions. **B)** IHC staining of 10 days old neuronal cultures with GFAP and MAP2 antibodies show the presence of neurons, but not astrocytes. **C)** IHC staining of 14 days old neuronal cultures with Homer1 and Basson antibodies show the formation of functional synapses. **D)** APOE2 and APOE4 particles were isolated from normal human APOE2/2 and APOE4/4 postmortem brains. Western blot analysis with APOE antibody shows the eluted APOE particles (left). Right side figure shows a gel with coomassie blue staining. Arrow marks indicate the appearance of similar bands in the film and coomassie-stained gel. **E-G)** Western blot analysis of proteins isolated from *PS19* neuronal culture, which were treated with APOEs for 4 days, shows that APOE4 treatment induced the AT8 levels, and APOE4-mediated tau pathology was diminished in the presence of APOE2. n=4, one-way ANOVA *P<0.05 and ****P<0.0001. IHC scale bars=20 µm.

**Supplementary figure 2.**
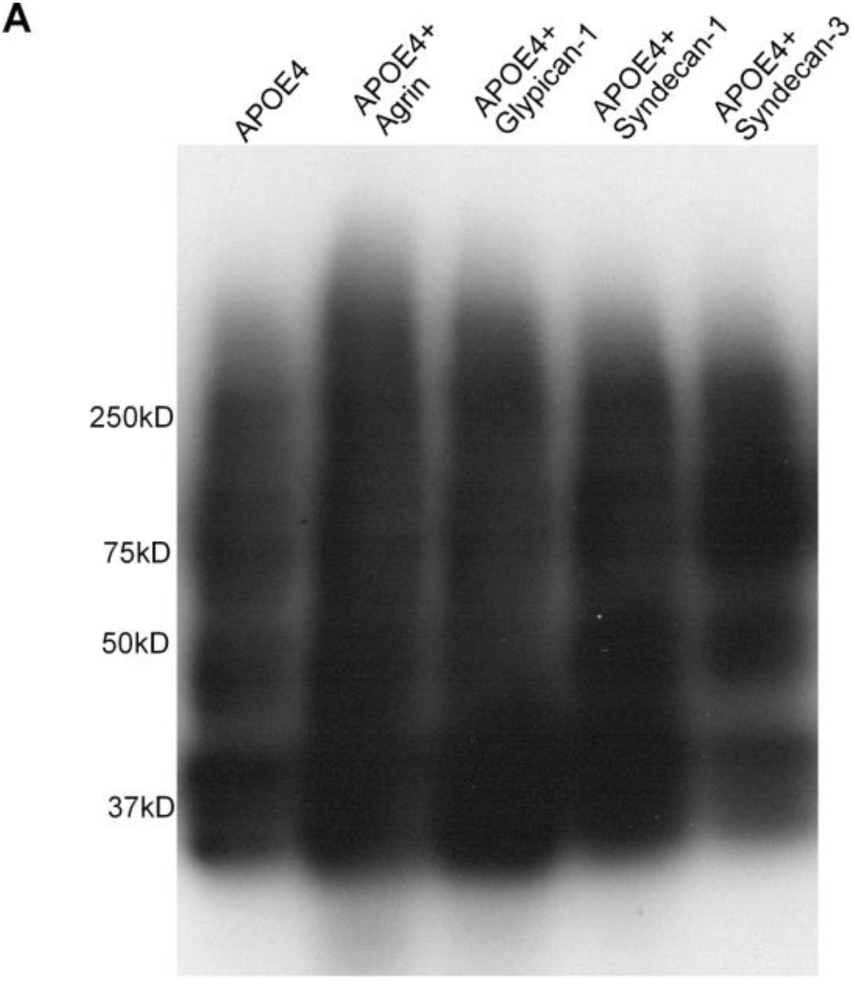
**A)** The HSPGs such as, Agrin, Glypican-1 and Syndecan-1 and Syndecan-3 proteins, were incubated with APOE4 at room temperature for 1h, and performed western blotting with anti-APOE antibody. No notable changes in the terms of protein shift were observed.

**Supplementary Figure 3.**
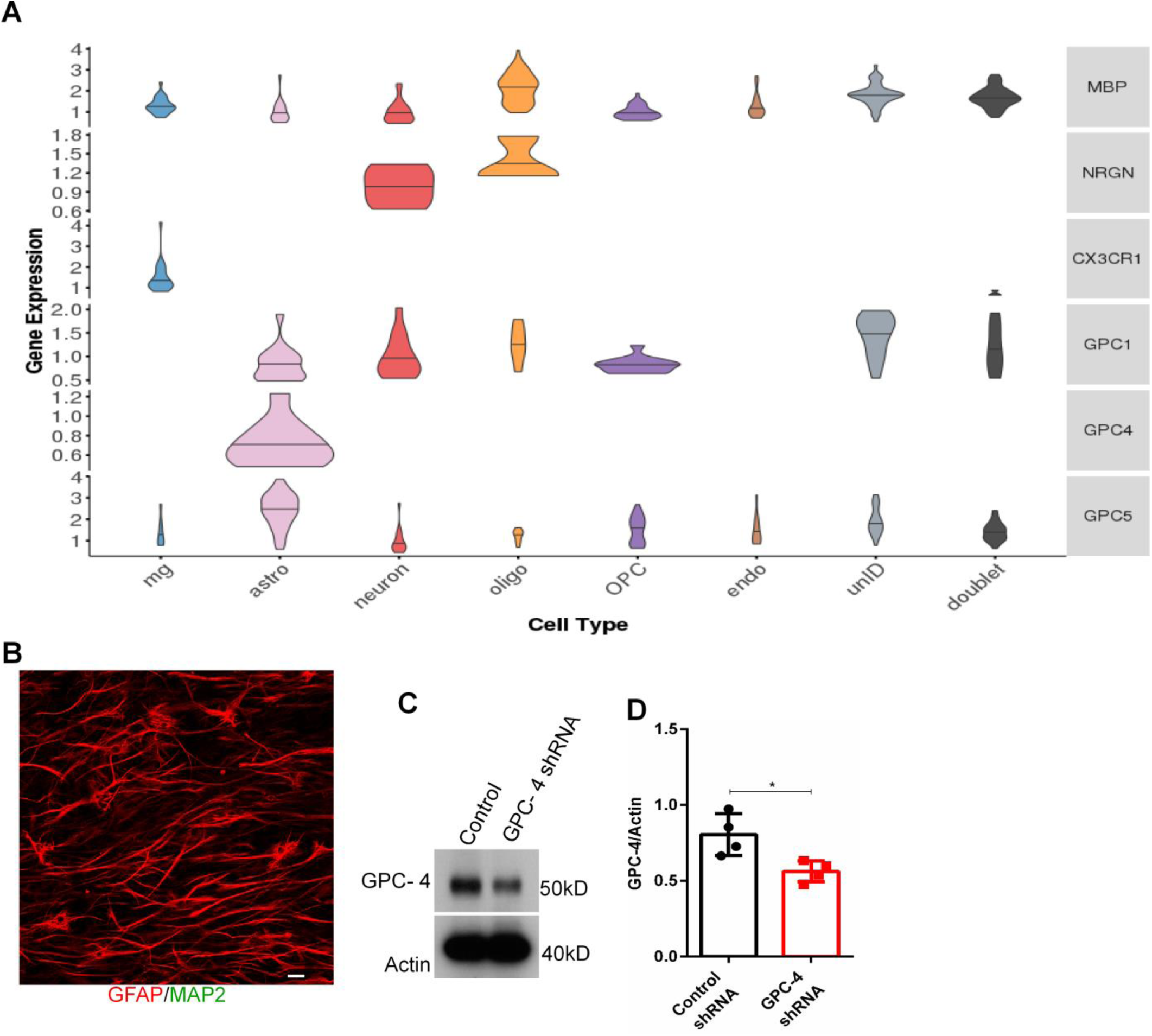
**A)** Violin blots from Grubman *et al, Nature Neuroscience*, 2019 show that GPC-4 is expressed in astrocytes. **B)** An IHC image of 12 days old astrocyte culture with MAP2 and GFAP antibodies show the presence of astrocytes, but not neurons. **C, D)** Western blot analysis shows that GPC-4 shRNA treated astrocytes expressed approximated 30-40% less GPC-4 proteins compared to control. n=4, unpaired t-test, *P<0.05. IHC scale bars=20 µm.

**Supplementary Figure 4.**
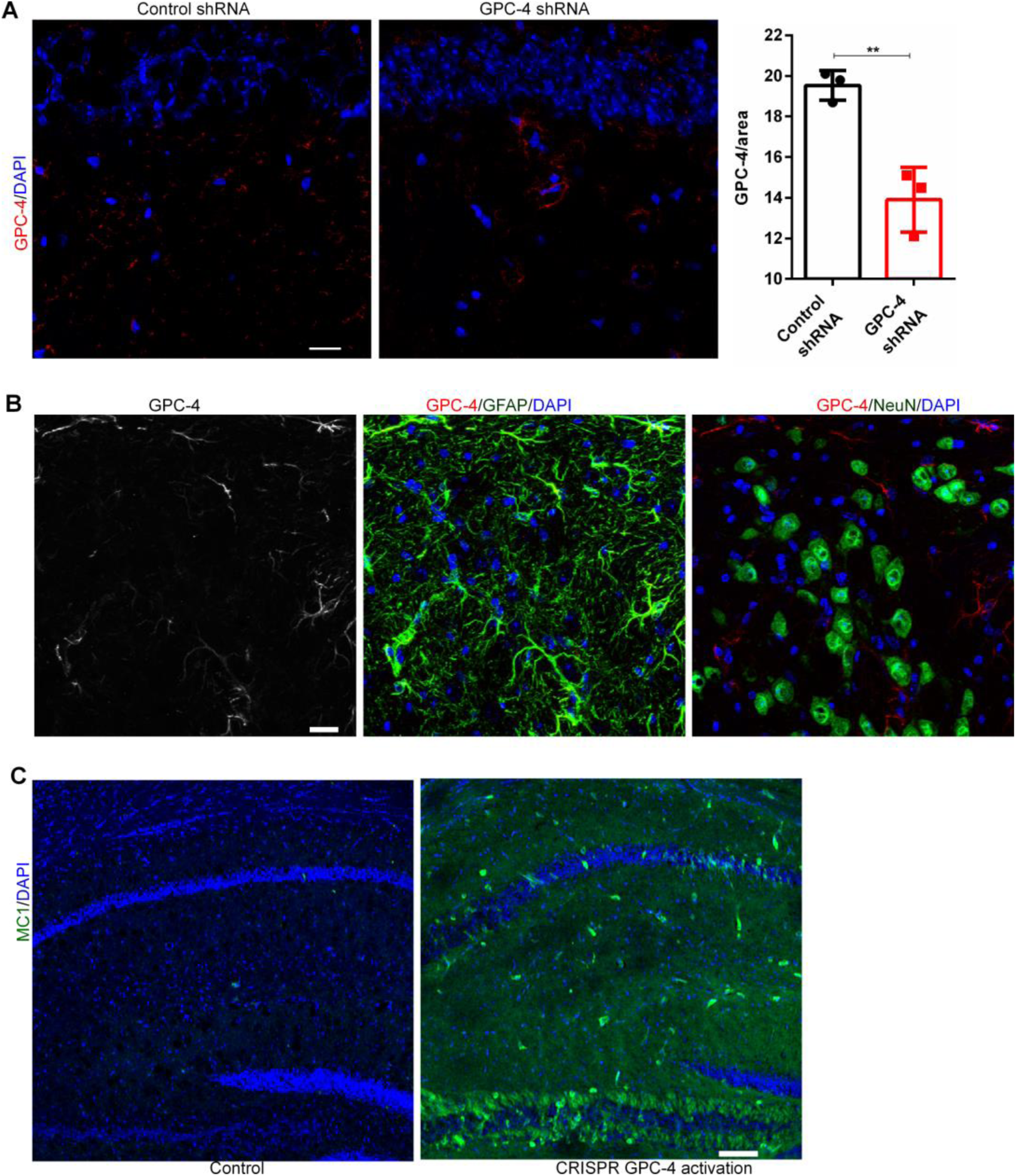
**A)** IHC staining with GPC-4 antibodies show that GPC-4 shRNA treatment reduced approximately 40% of GPC-4 expression. **B)** Representative IHC images with anti-GPC-4, GFAP and NeuN antibodies show that injection of CRISPR/dCas9 lentivirus GPC-4 activation system induced the expression of GPC-4 in astrocytes, but not in neurons. **C)** IHC staining with MC1 antibodies show that CRISPR/dCas9 mediated GPC-4 expression induced tau pathology *in vivo*. n=3, unpaired t-test, **P<0.01. IHC scale bars=20 µm and 100 µm.

**Supplementary Figure 5.**
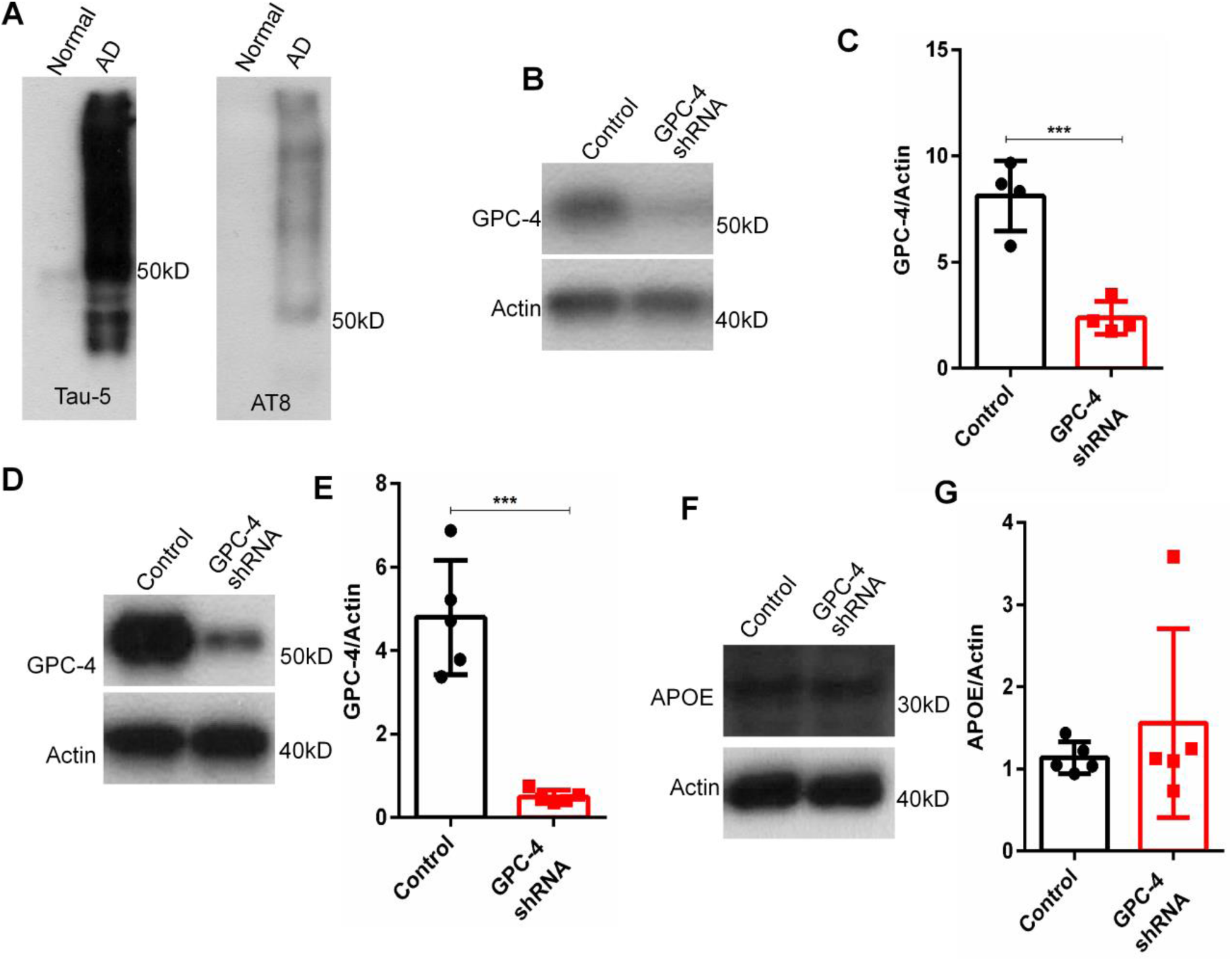
**A)** Western blot analysis with Tau-5 and AT8 antibodies show the isolation of insoluble tau proteins from postmortem AD brains. **B, C)** GPC-4 shRNA treated Tau FRET-biosensor cells expressed approximated 70% less GPC-4 proteins compared to control. **D, E)** Western blot analysis shows that GPC-4 shRNA treated iPSCs-astrocytes expressed approximated 80-90% less GPC-4 proteins compared to control. **F, G)** Western blot analysis shows that GPC-4 shRNA treated iPSCs-astrocytes expressed similar levels of APOE proteins compared to control. n=4-5, unpaired t-test, ***P<0.001.

**Supplementary Figure 6.**
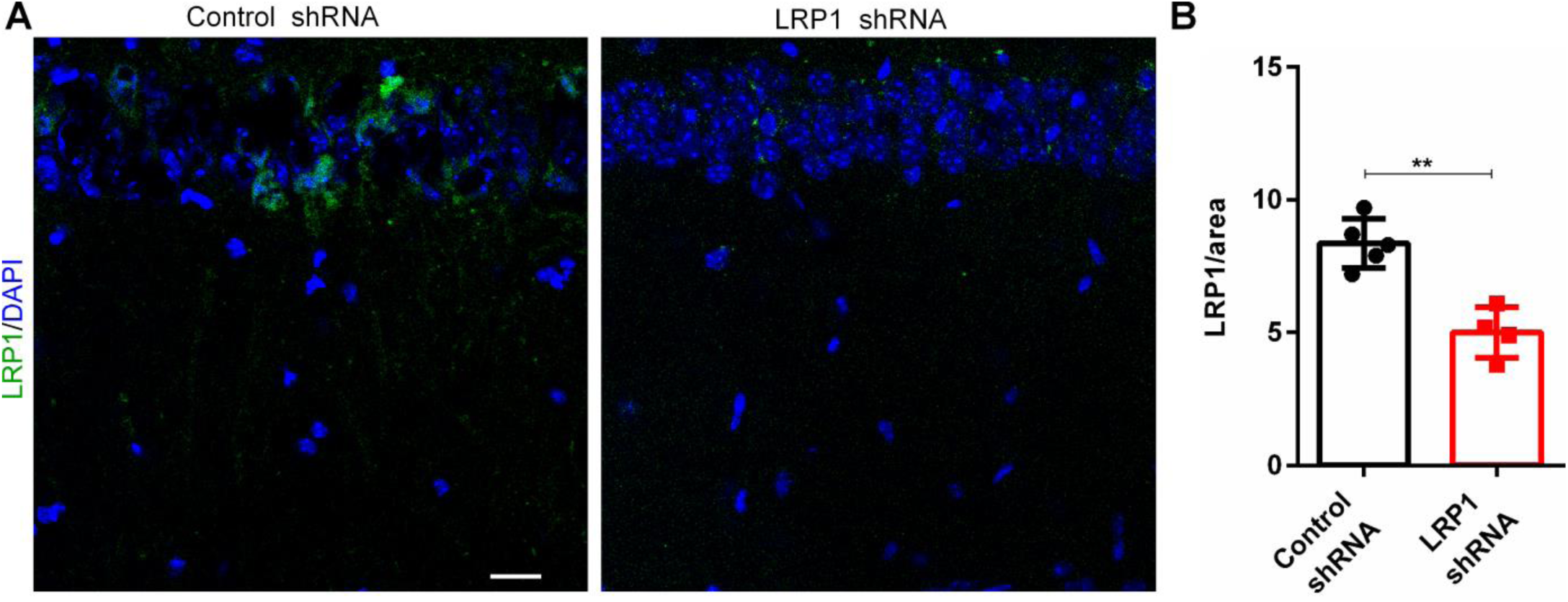
**A, B)** Representative IHC images with LRP1 antibodies show that LRP1 shRNA reduced approximated 40-50% of LRP1 levels compared to control shRNA. n=4-5, unpaired t-test, **P<0.01. IHC scale bars=20 µm.

## Materials and Methods

### Human postmortem brains

Human autopsy brain tissue from prefrontal cortical (PFC) regions (Brodmann area 10) were obtained from the Mount Sinai NIH NeuroBioBank and Banner Sun Health Research Institute. The sample details are given in the supplementary **table 1**. Pathologically confirmed AD cases were selected (Aβ deposits in neocortex and tau accumulation in the temporal /parietal cortical regions^65^). In addition, age-matched cognitively normal individuals were used as control in the experiments. Cases were not presorted based on the levels of tau or Aβ pathologies, race, sex and education.

### Astrocyte culture

The procedures were adapted from a previous publication^36^. Briefly, P1-2 mouse cortices were microdissected, digested with papain followed by trituration in low and high ovomucoid solutions. Cells were passed through a 20 µm mesh filter, resuspended in astrocyte growth media (DMEM supplemented with 10% FBS, Glutamax, sodium pyruvate, insulin and Penicillin/Streptomycin). The cells were incubated at 37°C in 5% CO_2_. Once it reached the 80% of confluences, cells were washed with PBS, and added conditioned medium^36^ and then incubated for three days. At the end of day 3, the medium was collected (ACM or astrocyte conditioned medium), concentrated/dialyzed using 10 kD Amicon^®^ Ultra Centrifugal filter tubes. The concentrated ACM was added to neuronal culture. For shRNA experiments, either GPC-4 shRNA or scrambled shRNA was added to astrocyte culture when incubated with conditioned medium to knock-down GPC-4. To study an interaction between microglial factors and GPC-4, astrocytes were treated with TNF-α (25ng/ml, #210-TA-005, R&D systems), IL-1β (25ng/ml, #201-LB-005, R&D systems) and IMD0354 (0.5 μM, #ab144823, abcam) for 2 days.

### Neuronal culture

Cortical neuronal culture was performed from *wild-type* or *PS19* or *MAPT*^*KO*^ P1 pups as indicated throughout the manuscript. The procedures were performed as described previously with minor modifications ^66^. Briefly, following dissection in Hybernate A solution, cortical tissues were digested with papain for 30 min. Papain solution was inactivated with ovomucoid inhibitor, tissues were dissociated, and passed through a 20-µm mesh filter. To remove the cell debris and immune cells, we performed negative immunopanning in Lectin 1 coated dishes with goat-anti mouse IgG+IgM and goat-anti rat IgG antibodies. A positive immunopanning with neural cell adhesion molecule L1 antibody was performed to enrich the neurons. Neurons were cultured in Neurobasal™ Plus Medium supplemented with B27, Glutamax and Penicillin/Streptomycin. A half-of medium was replaced every 3 days. 1) To investigate the role of APOE variants in tau uptake/spreading, we co-cultured the neurons from *PS19* and *MAPT*^*KO*^ at 1:10 ratio. Eleven days old neurons were treated with APOE2 particles (1μg/ml) or APOE4 particles (1μg/ml) or both (1μg/ml each) for 4 days (Fig. 1E, F). 2) To study an effect of GPC-4 on tau pathology, *PS19* neurons were treated on day 11 with 5 μg/ml of GPC-4 protein (#9195-GP-050, R & D systems) for 4 days (Fig. 3A-F). The same amount of GPC-4 proteins were used to treat *PS19* and *MAPT*^*KO*^ neuronal co-culture to investigate the effect of GPC-4 in tau uptake/spreading (Fig. 3O, P). 3) To study the role of APOE variants or GPC-4 on surface LRP1, WT neurons were treated for 24 h with APOE2 protein (5 μg/ml, #SRP4760, Sigma-Aldrich), APOE4 protein (5 μg/ml, #ab50243, abcam) or GPC4 (5 μg/ml) as described in figure 6. 4) To study the role of ACM on tau pathology, neurons were treated with ACM for 4 days as described before^36^.

### Isolation of surface proteins

To isolate cell surface proteins, WT neurons were cultured in 24-well plates as stated above. The cells from three wells/conditions were pooled together. Cell surface proteins were biotinylated and isolated using Pierce Cell Surface Protein Isolation Kit (#89881, Thermo Fisher). All the steps were performed at 4°C unless otherwise mentioned. The cells were washed briefly with PBS and incubated with Sulfo-SS-Biotin for 30 mins. After the reaction was quenched, the cells were centrifuged at 1,200rpm for 4mins, and the resulting pellet was lysed with a lysis buffer and incubated for 1 h. Some fraction of samples was collected at this stage to evaluate the total protein levels. After 1 h incubation with a lysis buffer, samples were centrifuged at 12,000 rpm for 5 mins. The supernatant was incubated with NeutrAvidin Agarose for 1 h in a rotating shaker. Unbound proteins were washed with a wash buffer 3 times. The slurry containing proteins was boiled for 5mins at 95°C with 50 mM DTT containing SDS sample buffer. Finally, the samples were centrifuged at 1,200 rpm for 2 mins to collect the proteins.

### Animals

All animal procedures were performed according to the Icahn School of Medicine at Mount Sinai Institutional Animal Care and Use Committee (IACUC) guidelines. 8 months old WT animals were used for tau propagation study using AAV eGFP-2PA-hTauP301L. We produced AAV eGFP-2PA-hTauP301L virus in collaboration with Penn Vector Core. 1μl of AAV eGFP-2PA-hTauP301L were injected in the CA1 regions (AP-2, ML-1.3 and DV-1.5) of the hippocampus and examined after 4 weeks. 4 months old *PS19* animals were used for the following experiments. To induce the expression of GPC-4, glypican-4 CRISPR/dCas9 lentiviral activation particles (sc-420640-LAC-2, Santa Cruz) were injected in the right CA1 region (AP-2, ML-1.3 and DV-1.5). And the same volume of control lentiviral activation particles (sc-437282, Santa Cruz) was injected in a separate cohort of animals for comparison. After one week of incubation period, animals were sacrificed and processed for IHC to access the expression of GPC-4. After 3 weeks of injection, the tau pathology was examined. In another study, 2μg of APOE4 or APOE2 particles were unilaterally injected in CA1 regions. After 24 h, one group received control shRNA lentiviral particles (sc-37007, Santa Cruz) + APOE2, one group of APOE4 injected animals received control shRNA lentiviral particles and another APOE4 injected group received GPC-4 shRNA lentiviral particles (sc-145457-V, Santa Cruz). The APOE particles and lentiviral particles were derived through the same injection site. Animals were sacrificed after 3 weeks of injection for further analysis.

### Protein binding assays

To investigate whether GPC-4 differentially interacts with APOE2 and APOE4, the protein binding assay was performed. The concentration of any indicated protein was at 1 μg/μl. The final volume was made to five microliters with 50mM Tris buffer. The mixture was incubated at a shaker for one hour. The samples were boiled at 95°C for 5mins with or without 2-mercaptoethanol, and separated by a native gel.

### Immunoprecipitation

The total protein was isolated from frozen human brain samples using 1% Triton in a 50mM Tris buffer. Pierce Direct IP kit (#26148, Thermo Fisher) was used to isolate proteins of interest. Briefly, the AminoLink Plus Coupling Resin was washed with a coupling buffer, and the resin was incubated with 10μg of LRP1 or APOE antibodies for 2 h in a rotator. The unbound antibodies were washed away, and 1-3 μg of isolated proteins was added to resin and incubated overnight, 4°C, in a rotator. It was washed three times to remove unbound proteins, and antibody-coupled proteins were eluted and verified with western blot. In addition, isolated APOE particles were also used for invitro/invivo studies. For this purpose, the isolated particles were dialyzed with sterile PBS to remove any contaminants.

### Electrophoresis and western blot

The samples were lysed with SDS-lysis buffer (Laemmli Sample Buffer, Biorad), boiled at 95°C for 5 mins with 50 mM DTT, and separated based on their molecular weights using TGX Stain-Free Gels (Biorad). The separated proteins were transferred to PVDF membrane, blocked with 2% BSA for 1 h, incubated with primary antibodies overnight at 4°C. Membranes were washed with PBST 6 x10 mins, incubated with HRP-conjugated secondary antibodies for 1 h, again washed with PBST 6 x10 mins, and then developed using ECL solution (Clarity Western ECL substrate, Biorad) and XRAY films (#NC1382078, Fisher Scientific Company). The following primary antibodies were used: TAU-5 (#AHB0042, Thermo Scientific), AT8 (#MN1020, Thermo Scientific), PHF1 (#PA5-101061, Thermo Scientific), Actin (#4970, Cell Signaling), LRP1 (#ab92544, Abcam), APOE (#ab183597, Abcam), GPC-4 (#13048-1-AP, Thermo Scientific, and #MABN528, Sigma-Aldrich).

### Tau uptake assay

On day 11 of neuronal culture, the cells were treated either with ACM with APOE4 or GPC-4 shRNA treated ACM with APOE4 for 24 h as described in the manuscript. Finally, the neurons were incubated with 1 μg/ml of human tau protein for 1h and washed with PBS 3 times before proceeding to immunohistochemistry.

### Isolation of tau seeds

Insoluble tau proteins from frozen postmortem AD brains were isolated using a Sarkosyl-detergent method as previously described^67^. Briefly, 10g of frontal cortical tissue was homogenized using a Dounce homogenizer in nine volumes (v/w) of high-salt buffer (10 mM Tris-HCl, pH 7.4, 0.8 M NaCl, 1 mM EDTA, and 2 mM dithiothreitol [DTT], with protease and phosphatase inhibitor) with 0.1% sarkosyl and 10% sucrose, and centrifuged at 10,000 g for 10 min at 4°C. Supernatants were collected and final sarkosyl concentration was made up to 1%. After 1h incubation at room temperature, the samples were centrifuged at 300,000 g for 60 min at 4°C. The resulted sarkosyl-insoluble pellets were washed once in PBS, sonicated (10 pulses at ∼0.5 s/pulse) followed by centrifugation at 100,000 g for 30 min at 4°C. The resulted pellets were resuspended in PBS, sonicated again (20∼0.5 s/pulse), and centrifuged at 10,000 g for 30 min at 4°C. From the final supernatants, tau proteins were measured with Tau-5 antibodies using ELISA according to manufacturer’s instructions (The Human Tau Solid-phase sandwich ELISA, #KHB0041, Thermo Scientific).

### Tau seeding assay

The Tau Biosensor (#ATCC CRL-3275) cells stably expressing the repeat domain of Tau conjugated yellow fluorescent protein (YFP) were cultured in 24-well coverslips at 37 °C, 5% CO2 in a complete medium [for 500ml: 450ml DMEM, 50ml FBS, 5ml Pen Strep (100x), 5ml glutamate (100x) and 5ml sodium pyruvate (100x)]. The cells were treated with shRNAs for two days, and then incubated with 0.6μg/ml tau proteins with/without 1μg/ml of APOE4 particles for 2 days. After two days of treatment, the cells were fixed with 4%PFA for one hour and YFP signals were imaged at SP5 Leica microscope. The number of tau inclusion containing cells were counted blindly and normalized with total number of cells (DAPI).

### Human *APOE* iPSC-astrocytes

The consent for reprogramming human somatic cells to hiPSC was carried out on hSCRO protocol 19-04 at Mount Sinai (J.TCW). CRISPR/Cas9 genome-edited isogenic *APOE* lines (TCW1 *APOE* 44 lines derived from an AD patient) were utilized for this study^47^. Human *APOE4*-iPSCs were differentiated into astrocytes as described before^47,48^. Once the cells reached about 80% of confluences, cells were washed with PBS, added conditioned medium with shRNAs, and incubated for three days. At the end of day 3, the medium was collected, concentrated/dialyzed using 10 kD Amicon^®^ Ultra Centrifugal filter tubes. The concentrated ACM was added to *PS19* neuronal culture and incubated for 4 days.

### Immunohistochemistry and imaging

1) The paraffin embedded human prefrontal cortical sections (10μm thickness) were deparaffinized in xylene and rehydrated in a graded series of ethanol (100%, 100%, 95% and 70%) and, then heat-induced antigen retrieval was performed with an antigen retrieval buffer (#ab93678, abcam). 2) Mice were deeply anesthetized and perfused with 100mM PBS, followed by 4% PFA in 100mM PBS. The harvested brains were post-fixed in 4% PFA in 100mM M PBS for 24hr followed by cryoprotection in a 30% sucrose solution for 48hr at 4 °C. 40μm thick serial coronal sections were cut on a cryostat. 3) The cultured cells containing coverslips were fixed for an hour with 4% paraformaldehyde.

After a brief wash with PBS, tissue slices/cells were incubated with 10% Donkey serum in PBST for 30 min. Primary antibodies were added and incubated for 48 h at 4°C, washed 3 × 10 mins with PBST, probed with corresponding secondary antibodies for 2 h, then washed again with PBST before counterstaining with DAPI. The tissues were mounted with Aqua-Poly/Mount and imaged using Leica TCS SP5 confocal laser scanning microscope. The identical imaging settings were applied for every set of experiments. The following primary antibodies were used: AT8 (1:500, #MN1020, Thermo Scientific), AceTau (1:500, #ITK0084, G-Biosciences), MC1 (1:500, a gift from Dr. Peter), MAP2 (1:500, #ab5392, abcam), LRP1 (1:200, #ab92544, abcam), HT-7 (1:200, #MN1000, Thermo Scientific), GPC-4 (1:200, # PA5-72360, 13048-1-AP, Thermo Scientific), GFAP (1:1,000, Aves Labs) and GFP (1:500, # ab290, abcam).

### Data analysis

Statistical analyses were performed with Graphpad prism software. Unpaired t-test or One-way ANOVA with Bonferroni correction were used. Quantification of western blots and immunohistochemical staining were performed using Image J software. a) The number of AT8 and aceTau positive cells were blindly counted in human brain slices. We used at least 3 slices/cases (Figure. 1A-C). b) We quantified intensity of AT8 and GFP/area using image J. To optimize the imaging conditions, we cultured *MAPT*^*KO*^ and *PS19* neurons separately and their respective AT8 and GFP signals were used as a baseline. The imaging area containing the *PS19* neurons alone was not included in quantification. (Figure. 1D-F). c) For western blot quantification, the intensity of each immunoreactive band was measured and then normalized to the corresponding beta-actin immunoreactive band. Normalized values were grouped and compared for statistical analysis. d) To analyze tau uptake (HT-7 staining), we imaged the neurons with 40x objective. Before staining, the coverslips were washed thoroughly with PBS to remove any extracellular tau. Four different places/coverslip were imaged. The intensity of HT-7 staining in the imaged area was quantified. (Figure. 5A, B). e) The number of AT8 and MC1 positive cells/areas were counted manually in a blinded manner (mouse tissues). 3-5 slices per animal were used for quantification purposes. f) Activated astrocytes were quantified as described before^32,40^. Briefly, astrocytes were imaged with 60x objective, z-series with 0.5μm distance. Number of cellular processes leaving the soma was assessed in the z-series. Based on the number of cellular processes at 15μm radius from the center of the cell body, the astrocytes were grouped into two categories as described in the main text. g) The intensity of GPC-4 per GFAP+ astrocyte was blindly measured with image J. Using 2D images, the presence of GPC-4 signals inside the astrocytes were confirmed and then their intensity was measured in stacked images. h) scRNAseq data was analyzed using a link provided by the authors^37^. The genes related to disease-associated astrocytes from a mouse scRNAseq study was retrieved^42^. These disease astrocytic genes were plotted along with the GPC-4 gene in human astrocytic subclusters^37^ to examine whether disease-associated human astrocytes express GPC-4.

